# Acute restraint stress impairs aversive memory retention but not memory formation

**DOI:** 10.1101/2025.07.25.666772

**Authors:** Aline Lima Dierschnabel, Diana Aline Nôga, Luiz Eduardo Mateus Brandão, Catherine Caldas de Mesquita, Diego de Aquino Câmara, Ywlliane da Silva Rodrigues Meurer, Felipe Porto Fiuza, Rovena Clara Galvão Januário Engelberth, Regina Helena da Silva, Jeferson Souza Cavalcante, Ramón Hypolito Lima

**Author notes:** Corresponding author: Post-Graduation program in neuroengineering, Santos Dumont Institute, Edmond and Lily Safra International Institute of Neuroscience, Macaíba, Rio Grande do Norte, Brazil. These authors contributed equally to this work (Aline Lima Dierschnabel, Diana Aline Nôga).

## Abstract

Stress can alter neurochemical signalling, affecting memory processing, but its underlying neurobiological mechanism remains unclear. Here, we investigate the effect of acute restraint stress (ARS) on long-term retention of aversive memory in rats.

We exposed the animals to either handling or ARS protocol and tested the rats in the plus-maze discriminative avoidance task (PMDAT). Also, we performed immunohistochemistry assays to unveil the effect of stress on neuronal activity.

We found that ARS immediately after training does not impair memory formation but hinders retention. Training triggers a peak of C-fos 1 hour later and a delayed 18-hour increase of Zif268 in the dorsal CA1. The same does not occur when ARS is experienced immediately after training.

We demonstrate the crucial role of Zif268 and C-fos signalling in maintaining PMDAT LTM. ARS is more relevant for memory retention than for memory formation of discriminative aversive memory.

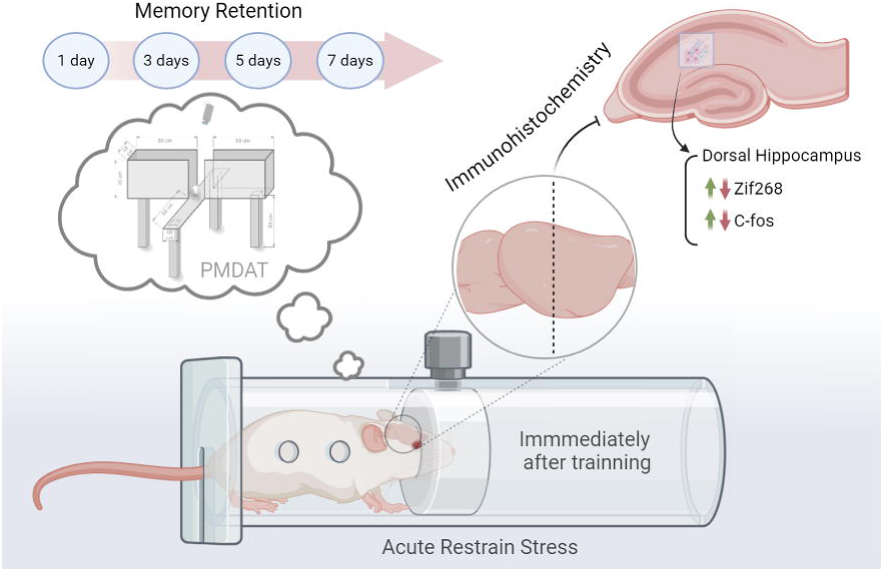

## 1. Introduction

Stressful events are an inherent part of life, with the potential to either promote resilience and personal growth or contribute to stress-related disorders. Although individuals may experience similar events, their responses vary, highlighting the diverse ways in which organisms adapt to adversity (Arnsten, 2015, Bittar et al., 2021).

The behavioural outcomes of stress exposure depend on how experience-dependent life events are acquired and stored. Stress can either impair or enhance memory function depending on various factors, including the nature of the stimuli, intensity, duration, controllability, timing within the memory processing phase, and the type of memory involved (Sandi & Haller, 2015).

Research findings on the effects of acute restraint stress (ARS) on memory are conflicting. Some studies report that ARS before learning disrupts mnemonic trace stability and impairs memory acquisition (Schwabe & Wolf, 2013; Shields et al., 2017; Beck et al., 2022; Wang et al., 2022), while others show enhanced learning, particularly for emotional content (Riggenbach et al., 2019; Vogel & Schwabe, 2016). Similar discrepancies exist for memory consolidation, with stress either strengthening or impairing it, depending on factors such as intensity and duration.

These opposing outcomes result from changes in neuronal activity within limbic regions, such as the hippocampus (Lupien & McEwen, 1997). Stress regulates cognitive functions and increases long-term memory (LTM) protein synthesis in the hippocampus (Espejo et al., 2017; Hebert et al., 2005; Lupien & McEwen, 1997; Sapolsky, 2002).

Dynamic neuronal gene expression, influenced by fluctuations in neuronal activity, supports neuronal plasticity. Hippocampal immediate-early genes (IEGs), such as C-fos and Zif268, activate and translate in neurons involved in learning processing (Guzowski et al., 2001). Evidence suggests that upregulation of C-fos and Zif268 expression in the dorsal hippocampus underlies LTM formation, consolidation, and storage (Katche et al., 2012; Revest et al., 2005; Veyrac et al., 2014; de Souza et al., 2019). The hippocampus is sensitive to stress, and dysregulation of C-fos expression in this region has been linked to stress-related disorders, including depression and anxiety (Schreiber et al., 1991; Miller & O’Callaghan, 2005).

We investigate the impact of post-training ARS on the maintenance of aversive memory and the expression of C-fos and Zif268 in rats subjected to the plus-maze discriminative avoidance task (PMDAT). This task assesses memory, anxiety, and locomotor activity (Silva & Frussa-Filho, 2000).

## 2. Materials and methods

### 2.1 Subjects and experimental procedures

Three-month-old male Wistar rats (280–300g, n = 128) were housed in home cages (4-5 animals per cage) under controlled conditions of temperature (22-25°C), humidity, with food/water at will and a light-dark cycle of 12h/12h (lights on 06:30 am). All animals were handled according to Brazilian law for the use of animals in scientific research (Law Number 11.794), and all procedures were approved by the local ethics committee (CEUA/UFRN - N° 039/2014). Animals were randomly assigned to each group.

### 2.2 Behavioural apparatus - Plus-maze discriminative avoidance task (PMDAT)

This apparatus consists of a modification of the elevated plus-maze (50 X 15 X 40 cm): two open arms (OA) opposite to two enclosed arms - a non-aversive enclosed arm (NAV) and an aversive enclosed arm (AV). In the training session, we placed animals in the centre of the maze, allowing them to explore the apparatus for 10 minutes. Every time animals entered the AV arm, an aversive stimulus (100W light plus 80-dB noise) was fired until the animal left the arm.

To determine memory retention in the PMDAT, animals were tested 1, 3, 5, and 7 days after training (n = 7-9 rats per group). Subsequently, we applied a 1-hour restraint stress immediately after training to determine stress’ influence on LTM memory retention. Animals were tested in the same time intervals as the previous experiment (n = 8-9 rats per group).

### 2.3 Acute Restraint Stress (ARS) Protocol

We exposed the animals to either handling or a 1-hour ARS protocol in a separate laboratory room. The apparatus consisted of an acrylic tube (6.5 × 19.8 cm) with ventilation holes to ensure proper breathing during exposure. Behavioural tests were conducted 24 hours after handling or ARS exposure. However, for immunohistochemistry experiments, animals were exposed to different conditions and euthanised at two time points, as described below. Animals were randomly assigned to each group.

### 2.4 Immunohistochemistry

To investigate the expression patterns of Zif268 and c-Fos following PMDAT, animals were euthanised and perfused at specific time points (n = 3–4 rats per group). The experimental groups included: (1) naïve animals, (2) naïve stress (rats exposed to the AV arm with light-noise pairings—five trials of a 3-second stimulus with a 5-second interval), (3) animals euthanized 1 hour after handling or (4) after ARS exposure, and (5) animals euthanized 18 hours after handling or (6) after ARS exposure.

After appropriate manipulation of each experimental group, rats were anaesthetised with thiopental (100 mg/kg) and transcardially perfused with phosphate-buffered saline (PBS), pH 7.4, followed by 4% paraformaldehyde in 0.1 M phosphate buffer, pH 7.4. Each brain was sectioned into 50µm slices with a cryostat microtome (Leica, Germany) at −22°C and placed sequentially in six series with an anti-freezing solution. The distance between one section and the next in the same series was approximately 300µm.

To detect Zif268 and C-fos proteins, we performed an ABC-free-floating immunohistochemistry protocol. Sections were incubated for 16-18h with anti-C-fos (Santa Cruz Biotechnology [sc-52], 1:1000) or anti-Zif268 (Santa Cruz Biotechnology [sc-189], 1:1000) primary antibodies.

The solution contained 1% albumin diluted in 0.4% Triton X-100 and 0.1 M phosphate buffer, pH 7.4, and then incubated with a biotinylated secondary anti-rabbit antibody (Jackson ImmunoResearch Laboratories Inc., 1:1000) for 2h. Slices were washed and incubated with avidin-biotin-peroxidase solution (ABC Elite kit, Vector Labs, Burlingame, USA) for 2 hours. The reaction was developed by adding diaminobenzidine tetrahydrochloride (Sigma, USA) and 0.3% H_2_O_2_ in 0.1 M phosphate buffer, pH 7.4. Sections were washed, dried, dehydrated in a graded alcohol series, cleared in xylene, and coverslipped with Entellan (Merck).

All immunostainings were performed concomitantly, minimising possible differences in background among animals. Sections were examined by using an optical microscope (Nikon Eclipse Ni-U) with a digital camera (Nikon, DS-Ri2) to record images.

To estimate the density of Zif268- and C-fos-positive cells in the dorsal hippocampus, we systematically selected four sections from each animal. Using the rat brain atlas (Paxinos & Watson, 2006), we identified brain sections located between −3.24 mm and −4.2 mm from the bregma. We analysed immunoreactive cells and measured hippocampal subregion areas with ImageJ software (version 1.48, NIH, USA).

### 2.5 Statistical analysis

Memory persistence in the PMDAT was evaluated by comparing total time (s) spent in the AV and NAV arms, and the percentage of total time spent in the AV arm [%TAV = time in AV / (time in NAV + AV) x 100]. Anxiety-like behaviour was evaluated by the percentage of time spent in open arms [% TOA = time in OA / (time in OA + NAV + AV) x 100] in training and test sessions. Locomotor activity was evaluated by total distance travelled (m) in training and test sessions.

We analysed the data using GraphPad Prism 9 with paired sample t-tests, one-sample t-tests, two-way ANOVA followed by Tukey’s post-hoc test, or one-way ANOVA followed by Tukey’s or Bonferroni’s post-hoc test, depending on the experimental requirements. We set statistical significance at p < 0.05.

## 3. Results

Our results indicate that animals submitted to the PMDAT responded to the aversive stimulus throughout the training session and preferred the NAV-enclosed arm (Figure 1B).

**Figure 1:**
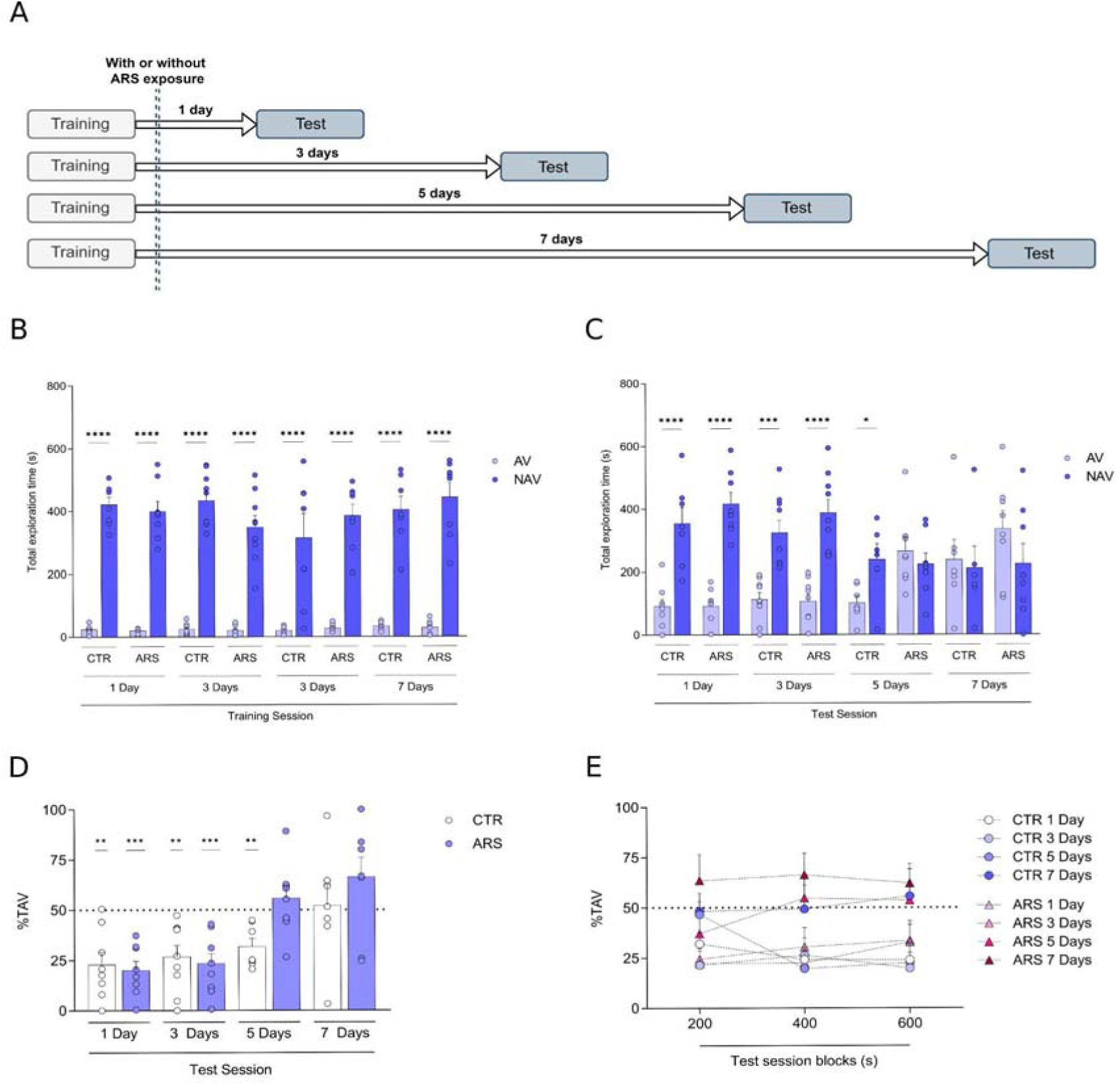
Memory retention in the plus-maze discriminative avoidance task (PMDAT). (A) Schematic diagram of experimental setup. Comparisons between total time spent in AV and NAV arms in (B) training and (C) test sessions. Percentage of total time spent in the aversive enclosed arm (%TAV) in the whole session (D) and in blocks of 200s (E). Data expressed as mean ± SEM. (B) * p<0.001 comparing AV vs NAV in all groups (paired sample t-test). (C) ** *p*<0.01 and *** *p*<0.001 comparing groups with a 50% chance of exploration of each arm (one-sample t-test). (D and E) All groups were compared among blocks (RM two-way ANOVA followed by Tukey’s *post-hoc* test).

When the percentage of time spent in the aversive enclosed arm was evaluated as a discriminatory proxy we found that control animals were able to discriminate enclosed arms for up to 5 days [Figure 1C; 1 day (t_(7)_ = 4.393; *p* < 0.001), 3 days (t_(8)_ = 4.098; *p* < 0.001) or 5 days (t_(6)_ = 4.544; *p* < 0.001)], however, we observed that ARS exposed rats showed an impairment of PMDAT memory retention compared to control group [Figure 1C; 1 day: t_(7)_ = 6.675; *p*<0.001; 3 days: t_(8)_ = 5.429; *p*<0.001; 5 days: t_(7)_ = 0.91; *p* =0.393; 7 days: t_(6)_ = 1.675; *p* = 0.1379].

When we analyzed the control group across time in the test session (blocks of 200s), the 7 days group showed a higher AV arm exploration in the last block when compared to all groups [Figure 1D; interaction - F_(6,54)_ = 3.017; blocks - F_(2,54)_ = 2.879; retention time - F_(3,27)_ = 3.175; followed by Tukey’s post hoc correction, * *p* < 0.05].

Moreover, we found that ARS animals showed a similar output when we analyzed the test session in blocks of 200s, however, 7 Days group is different from 1 and 3 Days in the first block and different than 1 Day group in the second block [Figure 1E; interaction - F_(6,62)_ = 1.153; blocks - F_(2,62)_ = 3.721; retention time - F_(3, 31)_ = 3.412; followed by Tukey’s post hoc correction, * *p* < 0.05].

We found changes in anxiety-like behavior [Figure 2A; F_(1, 56)_ = 5.128; p = 0.0274) and test (Figure 2C; F_(1, 56)_ = 4.317; *p*=0.0423)], and exploratory behavior comparing ARS groups to control groups during training [(Figure 2B; F_(1, 56)_ = 30.83; *p*<0.0001) and test (Figure 2D; F_(1, 56)_ = 17.44; *p* = 0.0001)].

**Figure 2:**
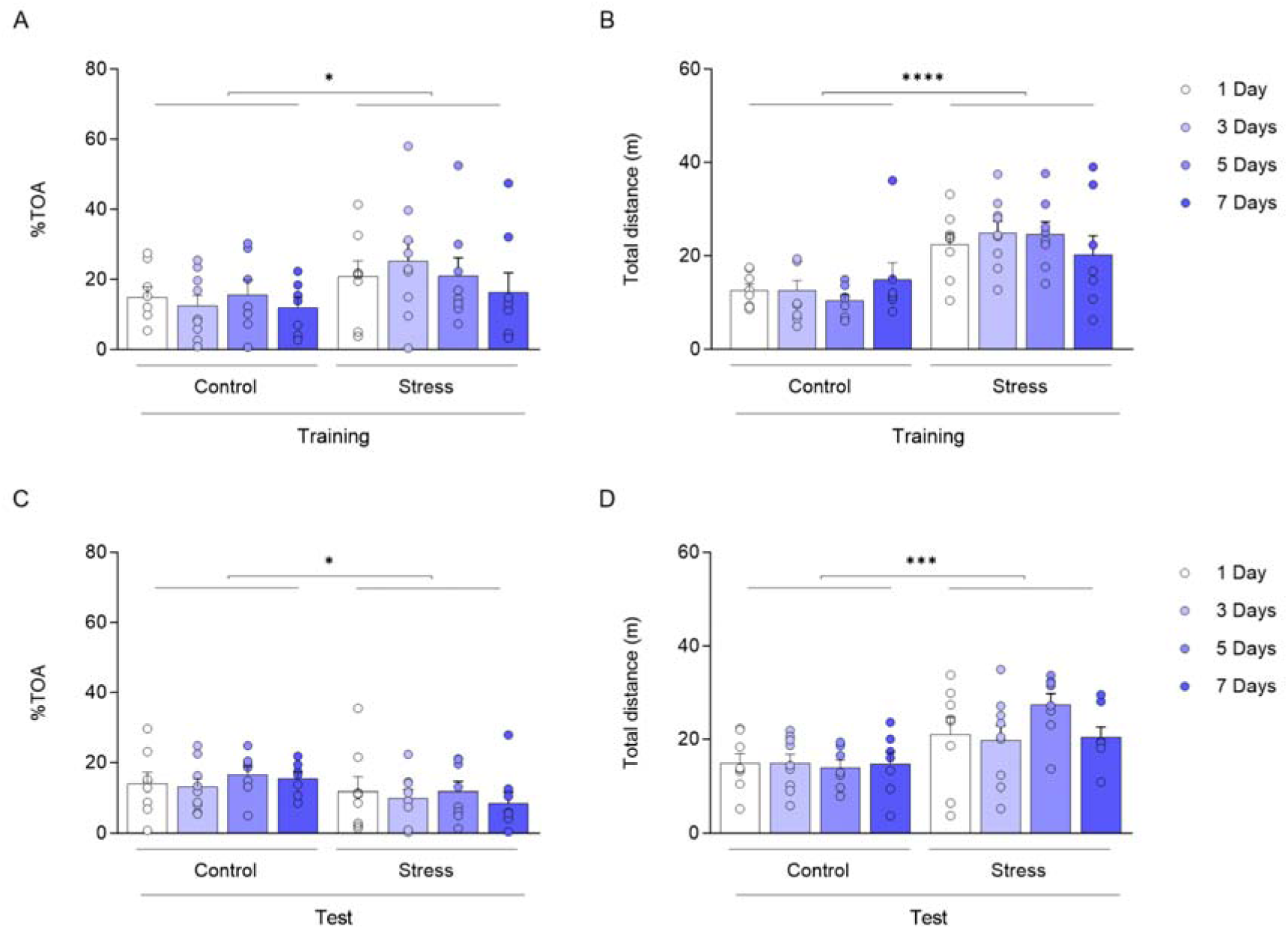
Anxiety-related behaviour and total exploration in PMDAT are not altered by training or ARS exposure. Percentage of time spent in open arms (%TOA) in the training session (A) and test session (C). Total distance travelled (m) in training session (B) and in the test session (D) for control and ARS animals, subsequently. Data expressed as mean ± SEM (Ordinary one-way ANOVA followed by Bonferroni’s post-hoc test).

Our results show that ARS exposure prevented the increase in the cell density of C-Fos at 1 hour [Figure 3; F_(5,17)_ = 7.713; *p* = 0.0006] and Zif268 at 18 hours [Figure 3; F_(5,18)_ = 9.580; *p* = 0.0001] after training in the CA1 subregion of the dorsal hippocampus. After multiple comparisons by Bonferroni’s post-hoc test, *p<0.05 and ***p<0.001 when compared to the naive group.

**Figure 3:**
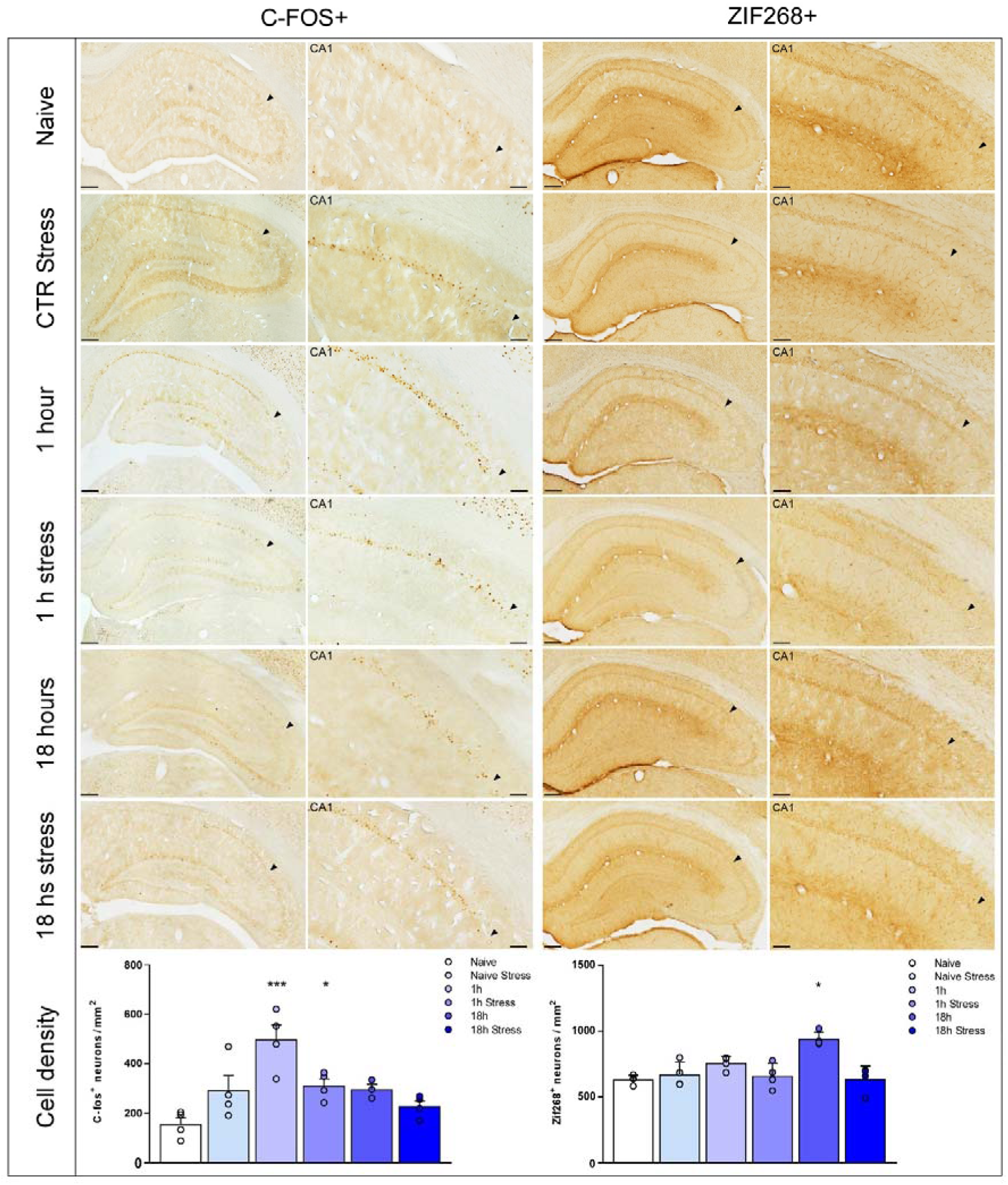
ARS immediately after training reverses C-Fos and Zif268 expression in the CA1 region of the dorsal hippocampus. C-fos (left) and Zif268 (right) immunoreactivities at the CA1 region of the dorsal hippocampus are indicated in each row for the distinct groups. The CA1 region is highlighted in higher magnification in the images on the right. Black arrows point to the limit between CA1 and CA2 hippocampal subregions. C-fos (Bottom left graph) and Zif268 (Bottom right) cell densities in the CA1 region of the dorsal hippocampus are shown at different time points after training in rats exposed to ARS and its respective control groups. Data expressed as mean ± SEM. Asterisks indicate the comparison with the naive group (one-way ANOVA followed by Bonferroni’s *post-hoc* test). Scale bar = 250 μm (dorsal hippocampus) and 100 μm (CA1 regions highlighted) in 100X magnification.

## 4. Discussion

The importance of our work relies upon understanding how acute stress modulates the hippocampal activity required to store information in long-term memories. We show that animals form and retrieve a persistent and reliable memory up to five days after training. This outcome allowed us to investigate the role of acute stress on the animal’s capacity to hold information about aversive memories. Other behavioural tasks can form stronger memories through electroshocks on rats’ paws as aversive stimuli (Lima et al., 2013; Rossato et al., 2009), but would jeopardise our ability to distinguish learning from stress induction.

PMDAT task requires both amygdala and dorsal hippocampus activity for memory formation and retrieval (Ribeiro et al., 2011; Leão et al., 2016). Nevertheless, the role of hippocampal subregions in the retention of PMDAT memory still needs to be understood.

The hippocampal subregions are often associated with different roles in memory formation and other facets. The CA3 subregion is thought to be involved in spatial memory and pattern separation (Gilbert & Kesner, 2006; Gilbert & Brushfield, 2009), while DG is associated with context discrimination (Ramirez et al., 2013). On the other hand, the CA1 subregion plays a crucial role in memory formation and persistence of LTMs (Neher et al., 2015; Rossato et al., 2009). It has also been suggested to be the primary output from the hippocampus to cortical areas (Amaral & Witter, 1989; Andersen et al., 2009).

Immediate-early genes (IEGs) regulate a wide range of genes and proteins and are often used to assess hippocampal activity levels (Marrone et al., 2011; Satvat et al., 2011; Vazdarjanova, 2004). C-fos and Zif268 have an essential role in the regulation of synaptic formation, transmission, and plasticity as well as memory processing (Abraham et al., 1991; Bozon et al., 2002; Gallo et al., 2018; Kim et al., 2018; Minatohara et al., 2016). IEGs upregulation in dorsal hippocampus pyramidal neurons underlies aversive memories’ long-lasting storage through MAPK/ERK pathway activation (Bekinschtein et al., 2007; Davis et al., 2003; Roux & Blenis, 2004; Veyrac et al., 2014). Multiple reports showed that C-fos and Zif268 inhibition impairs the retention of aversive memories by reducing BDNF expression (Bekinschtein et al., 2007; Katche et al., 2010), emphasising their relevance to this phenomenon.

C-fos and Zif268 are suggested to display a biphasic expression; an early peak, responsible for memory formation, and a late peak responsible for memory persistence (Bekinschtein et al., 2007; Katche et al., 2010; Katche et al., 2012). We found that Zif268 and C-fos cell density increased later after training in CA1. Contrariwise, only C-fos levels increased in other hippocampal regions, suggesting C-fos is sensitive to environmental stimuli such as novelty. Our results strengthen the idea that memory maintenance involves a late rather than early peak of Zif268 synthesis (Katche et al., 2012).

Acute stress promotes changes in hippocampal activity, impairing STM and LTM retrieval but enhancing LTM consolidation in humans and rodents. However, the effect varies depending on the nature of the events that trigger memory formation (Beckner et al., 2006; Buchanan et al., 2006; Buchanan & Tranel, 2008; Buss et al., 2004; Cahill & McGaugh, 1998; De Quervain, Roozendaal, & McGaugh, 1998; Diamond et al., 1992; Hui et al., 2006; Kuhlmann et al., 2005; Park et al., 2008; Preuß & Wolf, 2009; Smeets et al., 2008; Tollenaar et al., 2008). We found that ARS immediately after training did not alter memory formation, but reduced retention of PMDAT. Taken together, ARS differentially modulates LTM processing, hindering the persistence of LTM without affecting memory consolidation. However, neurochemical signalling and neural modifications underlying these behavioural changes are yet to be solved.

We repeated the neurochemical experiment to assess the effect of stress in either early (C-Fos) or late (Zif268) expression. We found that Zif268 cell density does not increase in non-stressed animals euthanised 18 hours after training. Moreover, stress did not cause any changes in animals euthanised 1 hour after training, suggesting that changes in Zif268 levels are related to retention of PMDAT memory. Conversely, we found the opposite results for C-Fos immunolabeling. Therewith, we did not find a stress effect *per se*, as naive animals submitted to stress induction did not show any disturbance in C-Fos and Zif268 levels.

Studies showed that ARS inhibits synaptic plasticity in the hippocampus-prefrontal cortex circuit in rodents (Rocher et al., 2004). Additionally, stress reduces the synthesis of other proteins for memory persistence, such as CREB, BDNF, C-Fos, and Zif268 (Espejo et al., 2017; Grønli et al., 2006; Kwon et al., 2007; Murakami et al., 2005). Collectively, one could argue that stress triggers an increase in corticosterone release in the hippocampus, modulating protein expression through its interaction with glucocorticoids and mineralocorticoids receptors, which could undermine LTM retention and long-term potentiation (LTP) maintenance (Jones et al., 2001; Veyrac et al., 2014).

Based on this rationale, increasing IEG synthesis in the hippocampus to promote the retention of PMDAT LTM (Antoine et al., 2014; Cheval et al., 2012) corroborates our results. Also, ARS immediately, but not late after training, decreases Zif268 expression in the same brain region (Gutierrez-Mecinas et al., 2011). In conclusion, we show the role of Zif268 and C-fos signalling in the formation and retention of PMDAT LTM. Both IEGs increase their expression in the hippocampus late after training. However, Zif268 overexpression has a high specificity in the dorsal hippocampus CA1 subregion, while C-fos increase its activity in

CA1 and CA3. We show that acute restraint stress immediately after training harms the maintenance of PMDAT memory and down-regulates Zif268 expression in CA1. Our results will help unveil the mechanisms underlying LTM maintenance under acute stress modulation. Persistence of an aversive memory and the stress situation surrounding learning play a key role in the development of psychiatric disorders, such as post-traumatic stress disorders. In other words, our results will benefit a broad range of studies, from basic research to clinical trials.

## 6. Author’s contribution

RHL and ALD conceptualised this work; ALD, DAN, LEMB, DAC, CCM, YRM, FPF, and RHL performed experiments, analysis, and artwork design; RHL, DAN, LEMB, and FPF wrote the original draft; RCGJE, RHS, JSC and RHL reviewed and edited the manuscript. RHS, JSC, and RHL funded this research.

## Acknowledgements

We thank Antônio Carlos Queiroz, Ezequiel Batista do Nascimento, Heloísa Lara Muniz, and Sarah Sophia Linhares for technical assistance.

